# Enhancing student comprehension of paternity assignment in molecular primatology: a pilot study using a Shiny web application in Kenya

**DOI:** 10.1101/2024.09.01.610690

**Authors:** David K. Mwaura, Jordan A. Anderson, Daniel M. Kiboi, Mercy Y. Akinyi, Jenny Tung

## Abstract

Kinship is a major determinant of affiliative and mating behavior in primates. In field studies, identifying kin typically relies in part on genetic analysis, especially for discriminating paternal relationships. Such analyses assume knowledge of Mendelian inheritance, genotyping technologies, and basic statistical inference. Consequently, they can be difficult for students to grasp, particularly through traditional lecture formats. Here, we investigate whether integrating an additional active learning approach—interaction with a Shiny web application*, DadApp*, which implements a popular paternity inference approach in an accessible graphical user interface—improves student understanding of genetic kinship analysis in molecular primatology. We do so in the context of a non-traditional learning environment in Kenya, a developing nation in which students have limited access to technology, and where the efficacy of educational Shiny apps has never been assessed. Twenty-eight (28) participants with diverse educational backgrounds attended an introductory lecture on genetics and paternity inference, completed a pre-test, interacted with *DadApp* via a structured set of exercises and questions, and then completed a post-test and survey about their experience and subjective understanding. Post-test scores significantly improved relative to pre-test scores (p-value=3.75 × 10^-6^ . Further, student interest and confidence in the subject matter significantly increased after the practical session with *DadApp*. Our results suggest that Shiny web app-based active learning approaches have potential benefits in communicating complex topics in molecular primatology, including in resource-limited settings where such methods have not yet experienced high penetrance.

**Graphical Abstract:** *Images in the graphical abstract are courtesy of publicdomainpictures.net

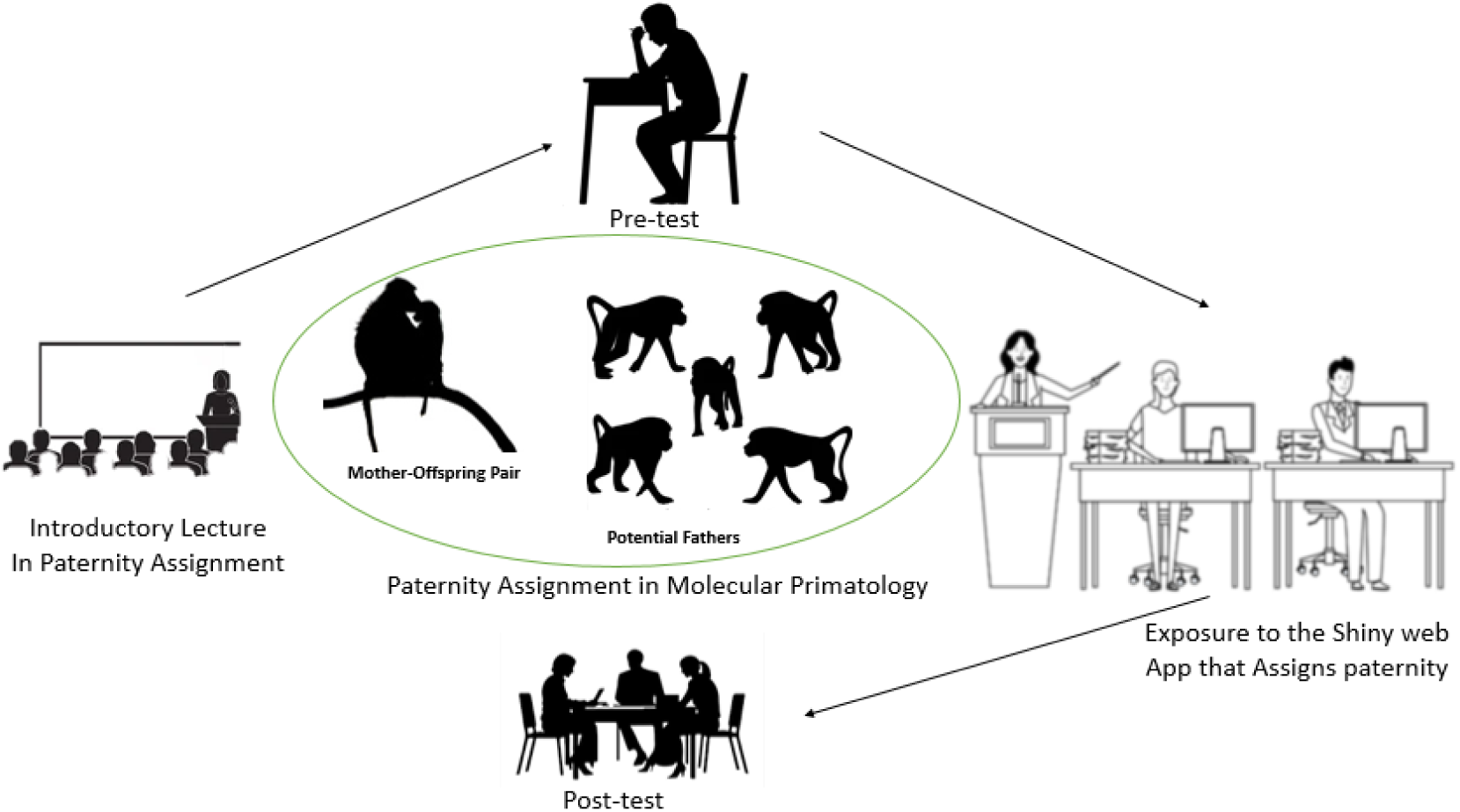

## Introduction

Kinship is one of the most important determinants of social organization, social structure, and mating patterns in primates (Strier, 2008; Chapais & Berman, 2004; Kappeler & Van Schaik, 2002). For example, in many primates, close kin preferentially affiliate with one another, but scrupulously avoid mating (Dal Pesco et al., 2021; Guo et al., 2015; Chapais, 2001). Indeed, avoiding mating with close kin is thought to be one of the most important selective factors governing the evolution of sex-biased dispersal, which characterizes approximately 75% of group-living primates (Morrison et al., 2023; Wellens et al., 2022; Galezo et al., 2022; Pusey, 1990). Selective investment in kin has also been demonstrated in many primate species. Both maternal and paternal kin preferentially associate with multiple species, including female rhesus macaques (Widdig et al., 2001), female baboons (Archie et al., 2014), female white-faced capuchins (Perry et al., 2008), male chimpanzees (Mitani et al., 2000), and male Assamese macaques (De Moor et al., 2020). Additionally, the identification of genetic fathers shows that males of even polygynandrous species can exhibit prolonged paternal investment (Städele et al., 2016; Buchan et al., 2003). For instance, baboon males maintain social bonds with the mothers of their previous offspring and intervene in agonistic interactions in favor of their juvenile offspring (Städele et al., 2021; Nguyen et al., 2009; Buchan et al., 2003). Together, these findings underscore the fundamental importance of identifying kin for studies of primate behavior and genetic structure.

While in some cases, kin relationships can be identified or reliably inferred through observational data alone, in most cases, at least some kin ties must be confirmed through genetic analysis (Städele & Vigilant, 2016). Studies in unhabituated animals, for instance, preclude observation of even close maternal kin (Masi et al., 2021). Additionally, even in highly habituated social groups, fertile females may mate with multiple partners in a conceptive cycle, leading to uncertainty about paternity and relationships among paternal kin (Rosenbaum & Silk, 2022; Platek & Shackelford, 2006). Consequently, many primate field studies infer relatedness by first using genetic data to reconstruct pedigrees that represent lineal descent, and then identify kin relationships based on the completed pedigree (as inferring non-parent-offspring kin classes directly from genotype data is typically quite noisy: (Van Horn et al., 2008)). Paternity (and/or maternity) inference is the crucial first step in this process.

Though methods for pedigree reconstruction and paternity inference in natural populations are well developed (Rentof et al., 2024; Petty et al., 2021; Sard et al., 2021), their application involves understanding several complex concepts. Students interested in pedigree inference in molecular primatology must, for instance, understand how alleles are transmitted through Mendelian inheritance. They also must understand how the accuracy of paternity assignment depends on both the quantity of genetic data available and the information content per genotyped locus, which in turn depends on genetic diversity in the species or population of interest. Additionally, students should understand how this information is used to then infer pedigree links. Doing so draws on concepts in both genetics and statistics, a combination that students can find challenging as they may not be covered together in one course or in relation to practical application in primatology (Fawcett & Higginson, 2012). Consequently, interested researchers or students may have only an abstract understanding of pedigree reconstruction using genetic data.

Such problems are not uncommon in ecology and evolutionary biology (Shou et al., 2015; Marquet et al., 2014). However, recent studies suggest that incorporating interactive learning tools can help students independently explore modelling and inference problems in biology and can help enhance student interest and confidence (Andrews et al., 2017; Haak et al., 2011; Nelson, 2008). For example, computer simulations can help improve student understanding of the mathematical underpinnings of population genetics (Hoban et al., 2012). Classroom exercises where students use simulated populations have also proven to be beneficial (Speth et al., 2010; Soderberg & Price, 2003; Winterer, 2001). In one study, for instance, active learning with population-level Punnett squares was shown to increase understanding and calculation proficiency in high-anxiety students, particularly for understanding Hardy-Weinberg equilibrium (Williams et al., 2021). Incorporation of games, clicker questions, and peer discussion or instruction can also help engage students more deeply in their learning materials (Subhash & Cudney, 2018; Smith & Wood, 2016; Kober, 2015; Vickrey et al., 2015). The common thread in these approaches is that they encourage students to become active participants rather than passive observers (Bernardo, 2017; Hague, 2011), allowing them to better understand and retain complex concepts in ecology and evolutionary biology.

One efficient and accessible method for developing active learning tools relies on the Shiny package (Kasprzak et al., 2021; Wang et al., 2021; Fawcett, 2018; Doi et al., 2016), an approach for web-based app development that builds on the popular R statistical programming language (Chang et al., 2015). R-based Shiny apps have become popular as teaching and data visualization tools in life sciences: as of 2022, more than 470 biological web applications have been developed and made accessible through online platforms such as shinyapps.io, CRAN, Github, or institutional websites and custom domains (Jia et al., 2022; Kasprzak et al., 2021). Their efficacy for instruction has been assessed in several ways. For example, González et al (2018) conducted a qualitative study where they evaluated the efficacy of Shiny apps based on student feedback. They concluded that students perceived Shiny apps as a useful educational tool for understanding probability, statistical inference, hypothesis testing, and modelling. Neyhard and Watkins (2020) also reported increased interest in and confidence with concepts in statistical genetics after interaction with Shiny simulations. Moore et al (2022) observed increased familiarity with ecological forecasting in undergraduate students after interacting with an interactive online module, built with R Shiny, which served as a visualization tool. These studies indicate that Shiny apps can provide useful practical experience for learning concepts in genetics, statistics, ecology, and evolution.

Shiny apps are accessible with minimal technological infrastructure: access to a computer, or, in some cases, a mobile device (Kasprzak et al., 2021). They also provide approaches for student interaction and inquiry that do not require wet lab facilities or reagents. These properties make them a particularly interesting avenue to consider in resource-limited settings, such as learning environments in developing nations. However, to date, studies that investigate the utility or efficacy of Shiny apps for life sciences education have focused on university settings in wealthy developed nations (e.g., Jia et al., 2022; Fawcett, 2018; Doi et al., 2016). Their potential as active learning tools for more diverse audiences, particularly in settings with more limited pedagogical and/or technological resources, has not been well-assessed.

To address this gap, we conducted a pilot study to test whether interaction with a Shiny web app, as a complement to a traditional lecture-based approach, improved student conceptual understanding of pedigree reconstruction in molecular primatology. Participants in the study were students and researchers with an interest in primate studies, but no specific background in genetic analysis or genetic paternity inference. First, we describe development of the app itself, *DadApp,* which implements the basic analytical algorithm in the program *CERVUS*, a standard tool for paternity inference in molecular ecology (Kalinowski et al., 2007; Slate et al., 2000; Marshall et al., 1998). *DadApp* allows users to provide biallelic genotype data and to calculate the relative log-odds (LOD scores) that a candidate individual is the parent of a given offspring of interest, given a set of possible candidates (Mwaura et al., 2023). We confirmed that *DadApp* correctly assigns paternity using data from a wild baboon population in the Amboseli region of Kenya (Alberts & Altmann, 2012). Second, we used a pre-test/post-test approach to assess the effectiveness of *DadApp* as an active learning tool outside of a university setting. Here, we gauged both its ability to increase student interest and engagement and its effect on improving conceptual understanding.

## Materials and Methods

### Application Development

We developed *DadApp* in R, one of the most common programming languages for biological data analysis and visualization (https://www.R-project.org/) (R Core Team, 2018), using the *shiny* package (Chang et al., 2017). *DadApp* is made up of two functional parts: ui.R, which specifies the user interface, and server.R, which specifies the calculations made on user-provided data and the resulting graphical representation of the results (Mwaura et al., 2023). To run, *DadApp* requires R version 3.4.0 or higher on a Microsoft Windows, Apple macOS, or Linux operating system, and can be run for data sets of up to 100 individuals and 200 loci using <1 Mb of storage space and <100 Mb of RAM. Upon execution, the application is launched in the user’s default web browser (e.g., Chrome, Firefox, Internet Explorer, or Safari), without requiring an internet connection.

*DadApp* operates under the simplifying assumption that the input data are biallelic genotypes, as in typical single nucleotide polymorphism data sets. Following convention, it represents homozygous reference genotypes as 0, heterozygous genotypes as 1, and homozygous alternate genotypes as 2. Following the approach described by Marshall et al., (1998), *DadApp* uses these data to determine locus-specific allele frequencies and then uses the rules of Mendelian inheritance to calculate the likelihood that a candidate parent is the true parent of a given focal offspring (i.e., the probability of the observed genotype data, under the hypothesis that the candidate is a true parent), relative to the likelihood that a random individual from the population is the true parent (i.e., the probability of the observed genotype data, given a model/hypothesis where the parent’s genotype is drawn randomly from the study population) (Kalinowski et al., 2010, 2007). This “background” likelihood is based on the allele frequencies in the population as a whole, which are calculated from the genotype data provided for the full, user-supplied data set. Values for each locus are combined across loci, under the assumption that each typed locus provides independent information (i.e., is unlinked).

If genotype data from a known parent are available, this information can also be provided to improve the accuracy of inference. Users can therefore provide these data by using the upload widget of the app. In primate studies, if one parent is known, it is typically the mother (Buchan et al., 2003). Therefore, in the scenarios we discuss below and *DadApp* itself, we refer to genotype data from the known parent as data from the mother, and treat the father as the unknown parent. However, the logic of the approach is identical if this scenario were reversed.

By rotating through the set of candidate fathers, each potential father is assigned a log-transformed likelihood ratio (LOD score). A positive LOD score indicates that the candidate’s genotype is more compatible with the genotype of the real father than a random genotype drawn from the population, while a negative LOD score indicates that the candidate’s genotype is less compatible with the genotype of the real father than a random genotype drawn from the population. The male with the highest LOD score is the algorithm’s top candidate. *CERVUS* provides an approach for assessing the confidence of top assignments (Slate et al., 2000; Marshall et al., 1998), but for simplicity here (and because the test data set used for the pedagogical analysis always included the true father), *DadApp* assigns paternity to the male with the highest LOD score. Following the approach in *CERVUS*, we also account for potential genotyping error by reassigning locus-specific likelihood values of 0, which can occur if a candidate father’s genotype is incompatible with the offspring’s genotype (Slate et al., 2000) (e.g., the father is CC but the offspring is TT), to a default value of 0.01. This approach avoids completely excluding candidate fathers based on a possible genotyping error, although incompatible values still considerably penalize the LOD score. This value can be changed by the user.

### Confirming accurate paternity assignments with pedigree and genetic data from wild baboons

To confirm that *DadApp* can make correct paternity assignments using genotypes and pedigree structures found in real populations, we drew on a subset of the data from Vilgalys et al., (2022), who analyzed whole genome resequencing data collected from a wild population of baboons in southern Kenya. This study population has been under continuous observation since 1971 (Alberts & Altmann, 2012), during which genetically assigned paternity has contributed to key insights about paternal care, reproductive skew, and inbreeding avoidance (Galezo et al., 2022; Alberts et al., 2006; Buchan et al., 2003). For the purposes of validating *DadApp* assignments, we subsampled local ancestry calls from 99 unlinked loci (this population is composed of hybrids between the yellow baboon, *Papio cynocephalus*, and the anubis baboon, *P. anubis*) and converted them to biallelic genotypes for all members of ten known pedigree trios (father, mother, and offspring). We then tested whether analyzing these data in *DadApp* resolved the correct father in each trio, within the full candidate set of all other males in the sample.

### Assessment of DadApp as an educational tool

Our primary goal was to assess whether an active learning component, here implemented using the *DadApp* interactive Shiny tool, improved student understanding of Mendelian inheritance, genetic data analysis, and paternity inference in the context of molecular primatology. To do so, we conducted a pre-test/post-test comparison of students and research staff at the Kenya Institute of Primate Research (KIPRE) in Nairobi, Kenya. KIPRE’s mission is to study nonhuman primates as models for human health and behavior. Most of KIPRE’s researchers do not primarily focus on genetics. Hence, participants in this study had a background in biology, but with limited formal training in genetics. Participants were recruited voluntarily into the study through direct outreach and referrals.

Participants (n=28) were given a one-hour introductory lecture on Mendelian inheritance, kinship, and genetic paternity. They then took an individual pre-test to assess their comprehension of these concepts (Supplementary Materials 1). After the pre-test, participants were asked to interact with *DadApp* to upload data, run analyses, and visualize results following a guided set of questions to explore alternative scenarios (e.g., data sets with or without maternal genotype data; data sets of differing size). The interaction phase took place in groups of 3-6 participants, 1-2 days after the lecture and pre-test. Subsequently, participants answered the pre-test questions again in a post-test phase. Notably, unlike in the pre-test phase, in the post-test phase they were able to engage in discussion with other group members. Finally, participants completed a post-test engagement survey in which they provided feedback on their understanding of key concepts before and after the active learning component (Supplementary Material 2). The survey included 5-level Likert-scale questions to quantify both perceived level of understanding (e.g. 1 -Not Confident, 2 -Somewhat Confident, 3 -Don’t Know, 4 -Confident, 5 -Extremely Confident) and interest/enthusiasm in the topic (e.g. 1 -Very Disinterested, 2 -Disinterested, 3 -Neutral/No opinion, 4 -Interested, 5 -Very Interested).

To assess the change in pre-test to post-test scores, we used paired Wilcoxon signed-ranks test to evaluate within-individual changes in performance following the active learning component. To evaluate whether this effect differed depending on characteristics of the student, we also used linear regression models to ask whether educational background, self-reported confidence, or self-reported interest predicted the percent change in pre-test to post-test score. In this study, we categorized educational background into three levels: undergraduates (current undergraduate students or students who had completed their undergraduate studies but no post-graduate education), MSc-level students (students currently enrolled in a Master’s program or students who had completed an MSc but no further education) and PhD-level students (students currently enrolled in a PhD program or who had completed a PhD). Finally, to assess whether the active learning component increased student interest in the topic, we analysed the feedback received from participants through their survey responses (Supplementary Material 3).

Ethical clearance for this work was granted through the Kenyatta National Hospital-University of Nairobi Ethics and Research Committee protocol no. P689/09/2023.

## Results

### Implementation of DadApp: an R Shiny package for exploring paternity inference and pedigree analysis

*DadApp* is freely available through Posit’s shinyapps.io platform at https://kiragu-mwaura.shinyapps.io/dadapp/, with additional annotated source code and instructions for installation and execution at https://github.com/KIRAGU-MWAURA/DadApp_Shiny_Web_App.

Examples of its user interface and outputs are shown in Figure 1, and example genotype data are available at https://github.com/KIRAGU-MWAURA/DadApp_Shiny_Web_App and the Supplementary Materials 4.

**Figure 1:**
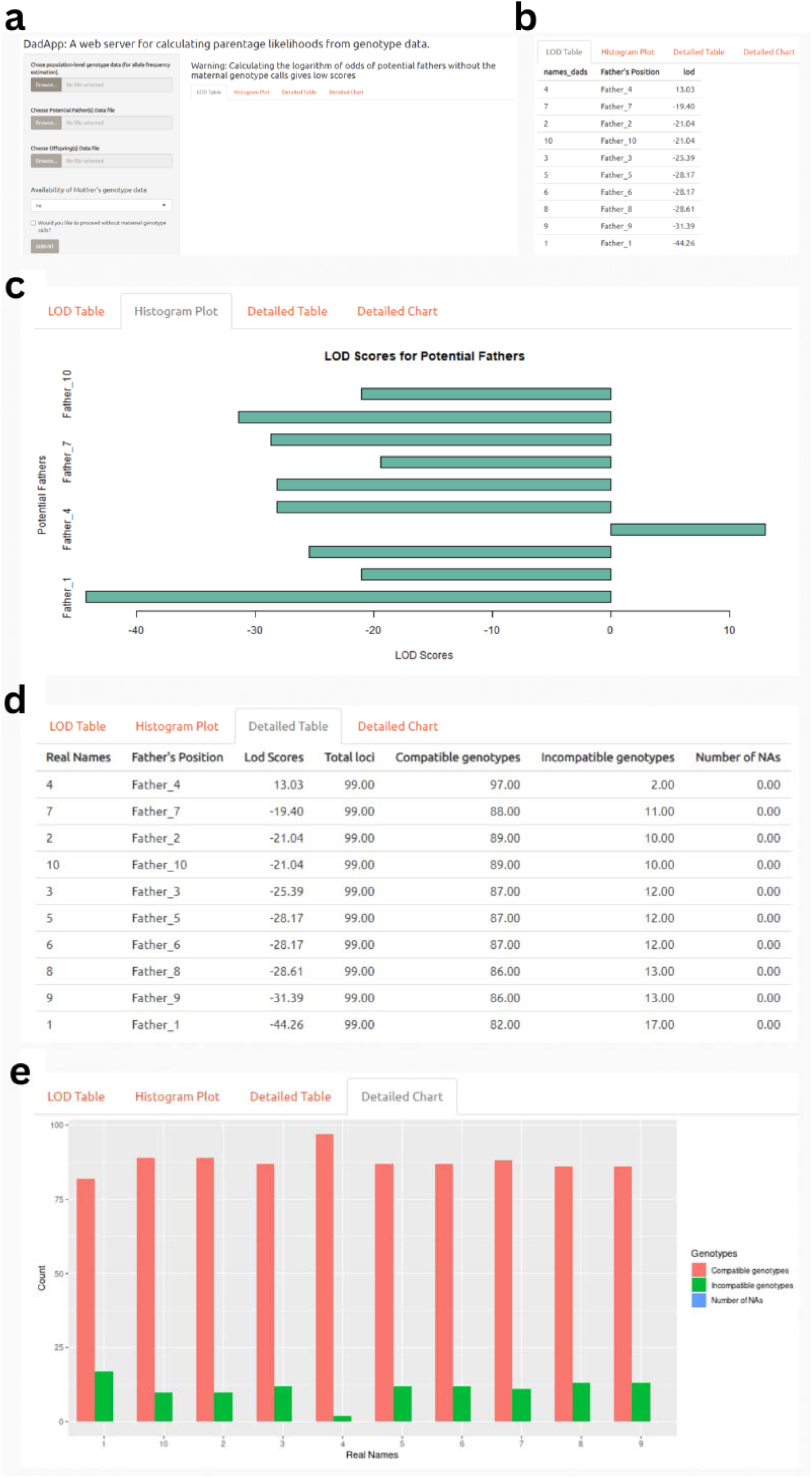
The *DadApp* user interface. a) The DadApp default user interface upon start up; b) Tabular representation of the app output, showing candidate fathers and their associated LOD scores; c) Barplot representation of candidate father LOD scores d) Detailed tabular representation of LOD scores for candidate fathers, including the total number of loci used to calculate the LOD score, number of compatible genotypes, number of incompatible genotypes, and number of loci with missing data; e) A barplot output showing the number of compatible genotypes, incompatible genotypes, and loci with missing data (denoted as NA) for each candidate father.

### *DadApp* correctly assigns paternity in trio data from wild baboons

We tested *DadApp* on genotype data from 99 biallelic loci, for 30 individuals (10 trios). In all cases, the highest LOD score candidate matched the independently assigned father in the Amboseli Baboon Research Project pedigree, where paternity assignment was based on genotyping data from 6 – 14 microsatellite loci (Alberts et al., 2006; Buchan et al., 2003). This result confirms that *DadApp* correctly implements the *CERVUS* algorithm in Shiny. It may therefore also be useful for basic paternity assignment in real data, provided the data types are compatible with *DadApp*’s capabilities.

### The utility of *DadApp* as an active learning tool

We recruited 28 volunteer participants into our study. Demographic data on self-reported gender, and educational level are shown in Figure 2.

**Figure 2:**
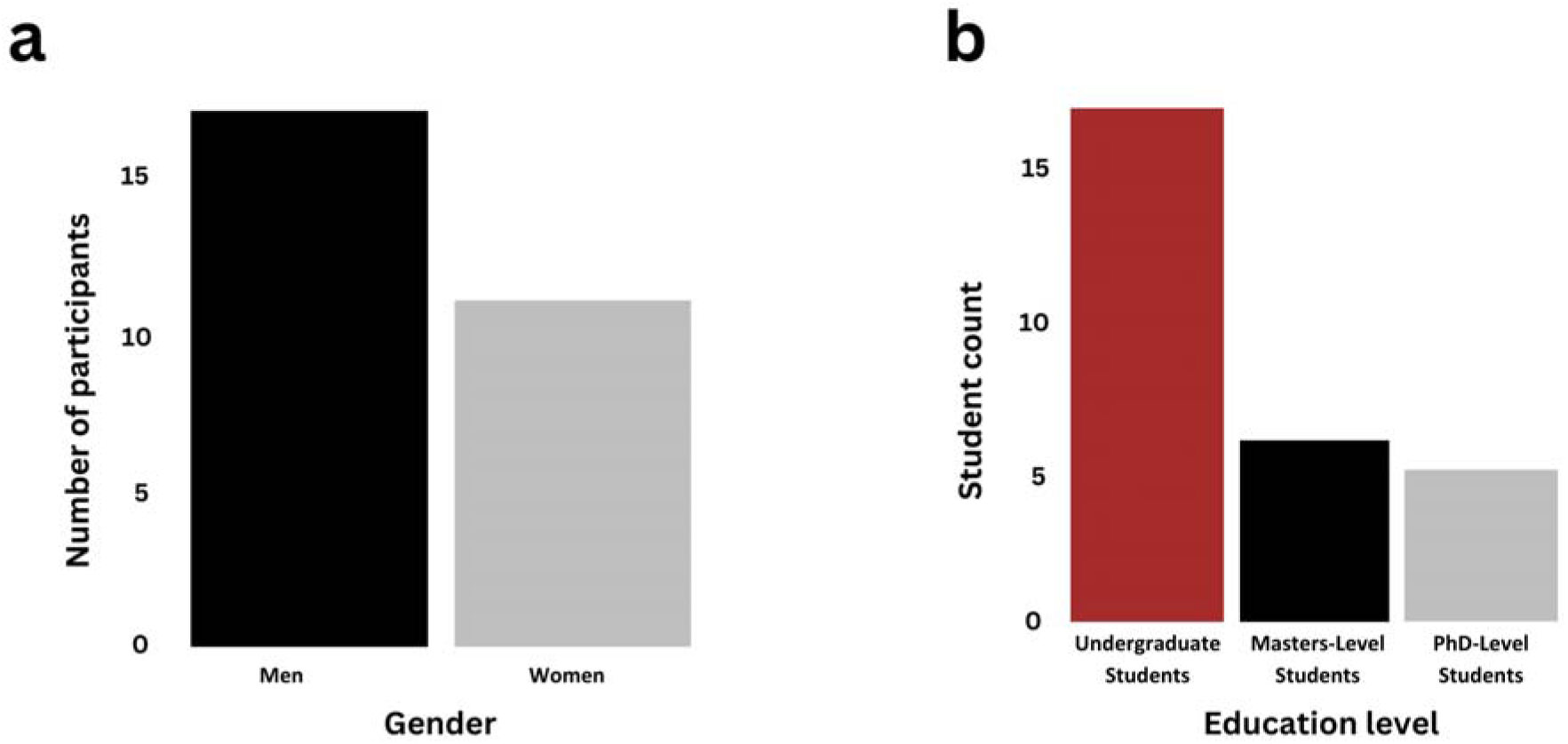
Participant information. Bar plot showing the distribution of participants by a) Self-identified gender and b) Self-reported education level.

The participants underwent tests before and after their interaction with the *DadApp* to evaluate their understanding of Mendelian inheritance, kinship, and genetic paternity inference. The mean score of participants across all subjects, out of a maximum of 100%, was 70.54% in the pre-test stage (standard deviation= 10.92, range= 45% - 90%) and 97.86% in the post-test stage (standard deviation= 4.18, range= 80% - 100%). We observed a statistically significant within-individual increase from the pre-test to post-test scores (paired Wilcoxon Signed-Rank test p = 3.75 x 10^-6^: Figure 3). Educational level did not affect test performance in either the pre-test (ANOVA: df =2, F-value =2.037, p-value =0.152) or post-test stages (df= 2, F-value= 0.584, p-value= 0.565). Self-reported confidence after the introductory lecture and self-reported interest at the beginning of the study did not significantly predict the percent change in the test score (linear regression: p-value= 0.652 and p-value= 0.9152, respectively).

**Figure 3.**
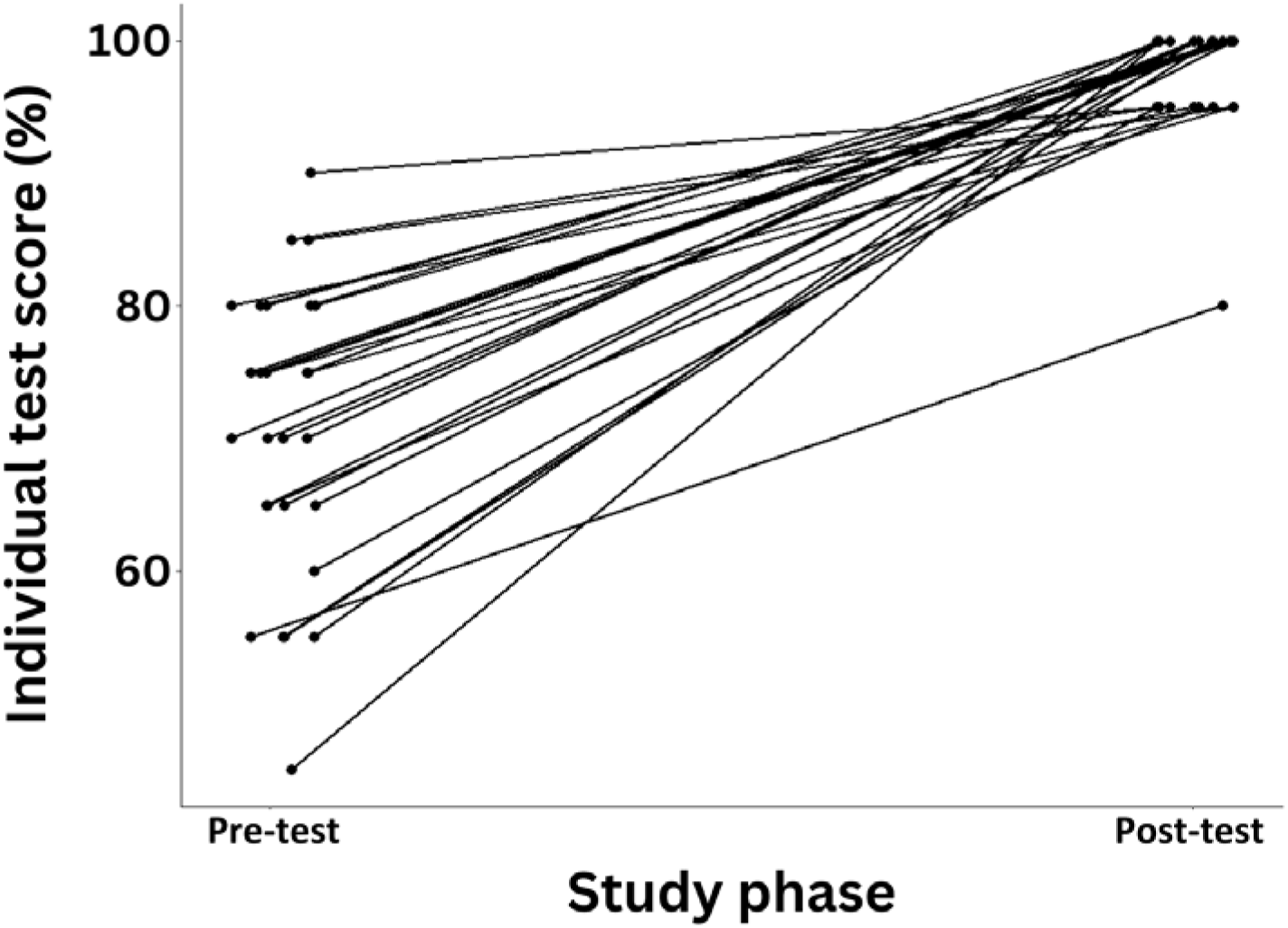
Within-individual comparison of pre-test and post-test scores after interacting with. DadApp. Lines connect the scores for the same individual between pre-test and post-test stages. The grouping of post-test scores may reflect discussion among individuals taking the post-test at the same time. Overall, post-test scores improved over pre-test scores (n=28, paired Wilcoxon Signed-Rank test p = 3.75 x 10^-6^). Points are jittered within pre-test and post-test values to avoid overplotting.

Finally, the participants completed a post-engagement retrospective survey to provide feedback on their experience. Participants’ ratings for most survey questions covered the entire 5-point scale. We observed no statistically significant difference in self-described interest based on the introductory lecture (mean rating provided retrospectively for interest before the lecture = 3.046 ± 1.48 s.d. versus interest after the lecture = 3.038 ± 1.40 s.d.; p-value=0.6114; paired t-test). Conversely, we observed a highly significant difference (p-value = 9.485 x 10^-5^; paired t-test) in self-described interest when comparing the post-introductory lecture responses to responses after the practical session with *DadApp* (mean = 4.746 ± 0.489 s.d.; Figure 4a). Similarly, we observed a statistically significant increase in participant confidence between the post-introductory lecture stage (mean=4.038 ± 0.958 s.d.) and after the practical session with *DadApp* (mean = 4.769 ± 0.815 s.d.; paired t-test p = 8.103 x 10^-4^) (Figure 4b).

**Figure 4:**
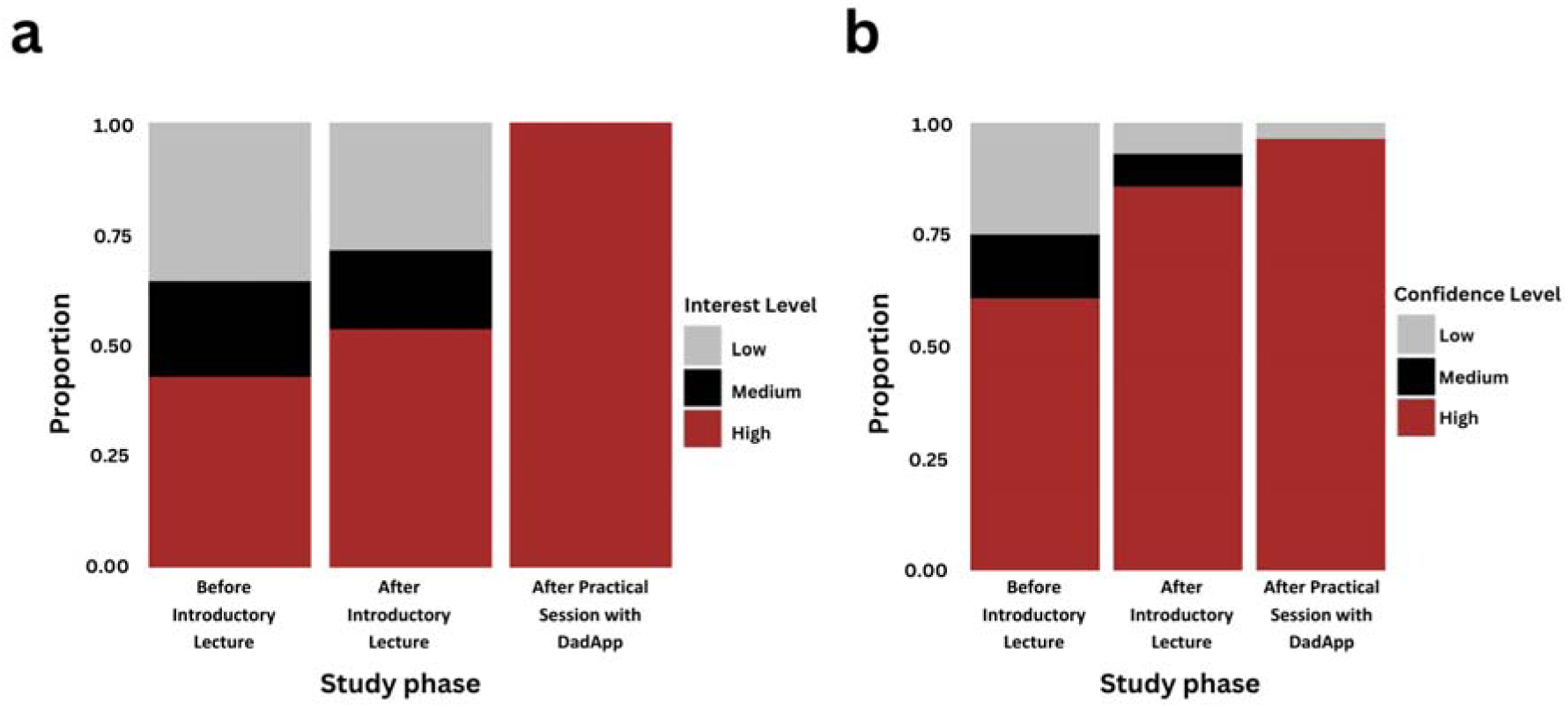
Participant interest and confidence throughout the study. . Bar plot showing a) Change in participant interest levels across stages of the study. b) Change in participant confidence levels across stages of the study

## Discussion

This study provides a first investigation of how Shiny apps can affect student learning and engagement in molecular primatology in a non-traditional classroom setting in the developing world. Our results show that Shiny apps can be effectively deployed in this context, where resources to support other forms of active learning (e.g., field courses, wet-lab practical) may be limited and non-traditional pedagogical approaches are not yet in wide use. And while we conducted this study at a suburban research institute, it is straightforward to envision using this or similar approaches even in field settings, where much of primate research is conducted in practice.

Exposure to *DadApp* as an instructional tool improved post-test performance compared to pre-test performance across educational levels. This result is consistent with work by Moore et al. (2020), who reported increased comprehension and interest in ecological modelling and forecasting for both undergraduate and graduate students when deploying an R Shiny application-based module. The use of technological enhancements and a collaborative learning environment has been repeatedly shown to improve student performance (Qureshi et al., 2023; Herrera-Pavo, 2021). Here, we generalize this finding, for the first time, to concepts in molecular primatology in a primate range country.

One of the most interesting results from our pilot is that, beyond test performance, student confidence and interest in kinship analysis and paternity inference increased significantly after interacting with *DadApp.* This finding also aligns with previous studies that reported enhanced engagement and student experience following exposure to interactive and application-based learning tools. For example, Neyhart & Watkins (2020) showed an encouraging increase in interest in statistical genetics after post-engagement with the *qshiny* Shiny app, which is designed to teach basic theory in quantitative and population genetics. Additionally, participant self-described confidence significantly improved after an active learning component in Moore et al. (2020), the R Shiny app-based ecological forecasting study. Overall, the literature suggests that interactive elements in educational software can foster deeper engagement and motivation among students (Clark & Mayer, 2016). Beyond comprehension, user experience and engagement are often linked to the interactivity, user-friendliness of the design, and relevance of the learning tool to real-world applications (González et al., 2018; Zhang, 2005).

Finally, while promising, we note some important limitations to the current study. The most significant limitation is that participants were allowed to take the post-test in small groups of 3-6, which likely inflated and reduced variance in the post-test scores and, consequently, the estimated effect of the app. Group dynamics, such as peer pressure, can lead participants to defer to the most confident or knowledgeable member of the group. Such dynamics can affect the accuracy and reliability of the post-test results as a direct assessment of the educational tool itself, as they can conflate individual-level with group- level comprehension. In retrospect, a more direct study design would have been to conduct the post-test in the same setting as the pre-test. An interesting possibility raised by our findings, though, is that peer discussion and interaction can amplify the effects of other active learning tools, as suggested by Freeman et al (2014). This possibility could be directly assessed in future work. A second limitation is that all assessments in this study were short- term. Consequently, we do not know how our deployment of *DadApp* affects long-term retention of the subject matter or participant interest. Finally, a potential downside of using educational apps is the significant amount of instructional effort and preparation required, including the time and resources needed for app design (Hanč et al., 2020; Doi et al., 2016). These costs can make active learning approaches more demanding than delivering a traditional lecture. However, it is worth noting that many developers are actively creating interactive biological web apps (Jia et al., 2022) and lesson plans, which may alleviate the burden on educators by reducing the need for new, extensive class preparation or development.

## Conclusions

Parentage analysis and pedigree reconstruction have become essential methods in molecular primatology. However, the concepts required to understand these methods can be difficult to teach. Our study suggests that incorporating active learning tools can help improve student understanding and, perhaps more importantly, student confidence and interest, even in non-traditional classroom settings in a developing nation context. Further, it shows that development of such tools using the Shiny platform is a feasible goal, and potentially another avenue for developing and learning skillsets important in modern primatology. Future studies should continue to evaluate the efficacy of these and other active learning approaches in diverse settings, with a diverse set of participants.

## Acknowledgments

We thank the Amboseli Baboon Research Project (ABRP) for financial support of the project. We also thank Arielle Fogel for providing local ancestry/genotype data for the Amboseli baboons to validate the Shiny App and all study subjects for their willingness to participate. The research in this study was approved by the Kenyatta National Hospital- University of Nairobi Ethics and Research Committee protocol no. P689/09/2023.

## Conflict of Interest

The authors declare no conflict of interest.

## Data Availability Statement

The data required to reproduce the analyses in this manuscript are provided as part of the Supplementary Materials.

## Notes

### Competing Interest Statement

The authors have declared no competing interest.

