## Supplementary Material 1 and 2 for "Enhancing student comprehension of paternity assignment in molecular primatology: a pilot study using a Shiny web application in Kenya"

**Supplementary Materials**

**Supplementary Material 1**

**PRE-TEST/POST-TEST QUESTIONS**

**Please read the following questions and answer appropriately.**

**Question 1: What role does statistical analysis play in paternity inference?**

a) It evaluates the physical resemblance between the alleged mother and child.

b) It measures the likelihood of paternity based on genetics.

c) It measures the emotional connection between the father and child.

d) It assesses the flow of genetic markers from offspring to parents.

Correct answer: c

**Question 2: What of the following statements is FALSE?**

a) Heterozygotes are individuals with dissimilar allele on homologous chromosomes
b) An organism with two identical alleles of a gene in their genome is called a homozygote
c) A recessive trait is a genetic characteristic that is expressed only when an individual carries two copies of the allele
d) Dominant traits are masked by recessive traits unless an individual inherits recessive allele from both parents

Correct answer: d

**Question 3: As minor allele frequencies (MAF) increase in a dataset, how might this impact the accuracy of genetic inference?**

a) Accuracy remains constant regardless of MAF

b) Accuracy increases

c) Identifies rare variants in the population which negatively impacts paternal inference

d) None of the above

Correct answer: c

**Question 4: If the frequency of the B allele in a finite population is given by p and the frequency of the alternative allele b is given by q, which of the following statements is false:**

a) If B and b are the only possible alleles at this locus, p = 1 – q
b) If two Bb heterozygotes mate, their offspring are expected to follow the ratio 1 BB: 2Bb : 1bb
c) Under random mating, the frequency of heterozygotes in the population is given by 2pq.
d) If B is selectively advantageous, b will definitely increase from one generation to the

next generation

Correct answer: d

**Question 5: What is the significance of having a shared genotype between an alleged father and a child in paternity testing?**

a) It increases the allele frequency of the shared genetic marker

b) It increases the probability of paternity

c) It decreases the likelihood ratio of the tested relationship

d) It indicates an error in the testing process

Correct answer: b

**Question 6: What information, in addition to genetic marker data, is ideally required for a more accurate paternity inference of an offspring?**

a) Only the genetic profiles of the alleged father and child

b) Genetic profiles of the mother, alleged father, and child

c) Only the genetic history of the alleged father

d) The allele frequency of the maternal lineage only

Correct answer: b

**Question 7:** **Why is it advisable to use multiple genetic markers rather than relying on a single marker for paternity testing?**

a) Single markers provide more accurate results

b) Multiple markers increase the complexity of the analysis

c) Allele frequency of single markers are the same in different population and geographical area

d) Multiple markers enhance the reliability and accuracy of results

Correct answer: d

**Question 8: Why is it important to consider the population's allele frequencies when performing paternity testing?**

a) Allele frequency determines the genotypic frequency of the offspring only

b) It helps identify potential mutations in both offspring and father's DNA

c) It provides a baseline for calculating likelihood ratios for patriarchal inference

d) It influences the child's height and weight

Correct answer: c

**Question 9: By itself, the equation: p + q = 1 tells us that**

a) The frequencies of alleles in a population are not changed by random mating

b) The sum of the frequencies of two alternate alleles at a locus must total 1.0

c) Genotype frequencies can be calculated if allele frequencies are known

d) There are only two possible genotypes in this population

e) A population that does not conform to this equation is evolving

Correct answer: b

**Question 10: You are a researcher about to embark on paternity inference. You calculate that you can afford to genotype 20 loci from a set of 40 known bi-allelic sites, in a fixed number of samples. To increase the power and accuracy of paternity assignment, you choose:**

a) The set of sites with the highest minor allele frequencies.

b) A set of 20 sites at random.

c) A set of sites that equally represent the range of minor allele frequencies in the possible set.

d) The set of sites with the lower minor allele frequencies.

Correct answer: b

**Highlight where the following statements are TRUE or FALSE**

**Question 11:** True or False: A high allele frequency indicates that this particular allele is less common in the population

Correct answer: False

**Question 12:** True or False: Calculating the likelihood ratio of potential fathers without maternal genotype calls results to low score of paternal inference

Correct answer: True

**Question 13:** True or False: Increase in amount of genetic information has negative impact on the statistical confidence on paternal assignment

Correct answer: False

**Question 14:** True or False: Allele frequencies is the same between different populations and geographical regions.

Correct answer: False

**Question 15:** True or False: Loci of rare variants are less informative in distinguishing individuals within a population.

Correct answer: True

**Question 16:** True or False: In paternity testing, the comparison of the child's genotype with those of both the mother and potential fathers helps determine the likelihood of biological parentage.

Correct answer: True

**Question 17:** True or False: Low allele frequency suggests that a particular allele is rare in the population.

Correct answer: True

**Question 18:** True or False: The accuracy of paternal assignment is positively influenced by the quantity of compatible genotypes shared between the alleged father and offspring.

Correct answer: True

**Question 19:** True or False: If both parents are homozygous for a specific locus, their offspring will also be homozygous for that locus.

Correct answer: True

**Question 20:** True or False: Allele frequency depends on the total number of alleles present, and multiple sets of genotype frequencies can be produced from the same allele frequency.

Correct answer: True

**Supplementary Material 2**

***This survey was originally administered through a Google Form.**

*DadApp Workshop Survey*

Title of Study:

Assessing *DadApp*, a Shiny Web Application that Assigns Paternity in Wild Animal Population, as an Educational Tool in Ecology.

The purpose of this e-survey is to find out if *DadApp*, a Shiny web application that implements commonly used pedigree inference and paternity assignment tools, improves user understanding of Mendelian inheritance, genetic kinship and relatedness, and paternity inference is a useful active learning tool for teaching these concepts .Participants in this research study will be asked questions about Mendelian inheritance, genetic kinship and relatedness, and paternity inference. Your response are anonymous.

This study is being conducted by Dr. Mercy Akinyi, Mr. David Kiragu, Dr. Daniel Kiboi and Prof. Jenny Tung. By completing the Pre/Post surveys, you express your consent to participate in the study. Thank you.

* Indicates required question

1. Your gender?*

*Mark only one oval.*

Male

Female

Prefer not to say

2. Level of Education?*

*Mark only one oval.*

Undergraduate

Graduate

Master's student

PhD student

Post-Doctorate

Please rate your level of interest in the following subjects in ecology before you went through an initial introductory lecture, after the lecture and after the *DadApp* practical?

3. Allele frequency calculation in animal population?*

*Mark only one oval*

Very Disinterested

Disinterested

No Opinion/Neutral

Interested

Very Interested

4. Hardy Weinberg Equilibrium?*

*Mark only one oval*

Very Disinterested

Disinterested

No Opinion/Neutral

Interested

Very Interested

5. Gene Flow in wild animal population?*

*Mark only one oval*

Very Disinterested

Disinterested

No Opinion/Neutral

Interested

Very Interested

6. Genetic variation in parentage analysis?*

*Mark only one oval*

Very Disinterested

Disinterested

No Opinion/ Neutral

Interested

Very Interested

7. Heritability in wild animal population?*

*Mark only one oval*

Very Disinterested

Disinterested

No Opinion/Neutral

Interested

Very Interested

As a result of the introductory lecture, what is your level of confidence in your knowledge of the following subject of paternity as a measure of ecological monitoring in wild animal population.

8. What is a test cross, and why is it used in Mendelian genetics?*

*Mark only one oval*

Not Confident

Somewhat Confident

Don't Know

Confident

Extremely Confident

9. Determining possible genotypes of offsprings' and their associated probabilities of occurrence given parents' genotype?*

*Mark only one oval*

Not Confident

Somewhat Confident

Don't Know

Confident

Extremely Confident

10. The concept of the likelihood ratio computational algorithm in the context of determining paternal inference*

*Mark only one oval*

Not Confident

Somewhat Confident

Don't Know

Confident

Extremely Confident

11. The role genetic markers play in the likelihood ratio computational algorithm for paternal inference*

*Mark only one oval*

Not Confident

Somewhat Confident

Don't Know

Confident

Extremely Confident

12. Calculation of allele frequency of bi-allelic markers in a wild animal population.*

*Mark only one oval*

Not Confident

Somewhat Confident

Don't Know

Confident

Extremely Confident

13. Steps involved in calculating the likelihood ratio for a single genetic marker in the likelihood ratio computational algorithm*

*Mark only one oval*

Not Confident

Somewhat Confident

Don't Know

Confident

Extremely Confident

14. How are the likelihood ratios of multiple genetic markers combined to obtain an overall combined likelihood ratio in the likelihood ratio computational algorithm*

*Mark only one oval*

Not Confident

Somewhat Confident

Don't Know

Confident

Extremely Confident

As a result of the practical session using DadApp, how useful/not useful are the following concepts useful when determining and/or assigning paternity in wild animal population.

15. Understanding the concept of statistical genetics for example probability using likelihood ratio*

*Mark only one oval*

Not at all Useful

Somewhat Useful

Moderately Useful

Very Useful

Don't Know

16. Genetic markers, Genetic variation and Gene flow*

*Mark only one oval*

Not at all Useful

Somewhat Useful

Moderately Useful

Very Useful

Don't Know

17. Allele frequency calculation as a factor in the concept of likelihood ratio computational algorithm*

*Mark only one oval*

Not at all Useful

Somewhat Useful

Moderately Useful

Very Useful

Don't Know

18. The importance of test cross in Mendelian genetics and heritability*

*Mark only one oval*

Not at all Useful

Somewhat Useful

Moderately Useful

Very Useful

Don't Know

19. Utilization of Hardy Weinberg Equilibrium in likelihood ratio computational algorithm*

*Mark only one oval*

Not at all Useful

Somewhat Useful

Moderately Useful

Very Useful

Don't Know

Please indicate your level of agreement with the following statements before the introductory lecture, after the introductory lecture and after the practical session using DadApp.

20. I have a better understanding of paternity inference using likelihood ratio algorithm in assigning LOD scores to putative fathers

Before Introductory Lecture: *

*Mark only one oval*

Strongly disagree

Disagree

Don't Know

Agree

Strongly agree

21. After Introductory Lecture: *

*Mark only one oval*

Strongly disagree

Disagree

Don't Know

Agree

Strongly Agree

22. After Practical Session: *

*Mark only one oval*

Strongly disagree

Disagree

Don't Know

Agree

Strongly Agree
